# Massive Mining of Publicly Available RNA-seq Data from Human and Mouse

**DOI:** 10.1101/189092

**Authors:** Alexander Lachmann, Denis Torre, Alexandra B. Keenan, Kathleen M. Jagodnik, Hyojin J. Lee, Lily Wang, Moshe C. Silverstein, Avi Ma’ayan

## Abstract

RNA-sequencing (RNA-seq) is currently the leading technology for genome-wide transcript quantification. While the volume of RNA-seq data is rapidly increasing, the currently publicly available RNA-seq data is provided mostly in raw form, with small portions processed non- uniformly. This is mainly because the computational demand, particularly for the alignment step, is a significant barrier for global and integrative retrospective analyses. To address this challenge, we developed all RNA-seq and ChIP-seq sample and signature search (ARCHS4), a web resource that makes the majority of previously published RNA-seq data from human and mouse freely available at the gene count level. Such uniformly processed data enables easy integration for downstream analyses. For developing the ARCHS4 resource, all available FASTQ files from RNA-seq experiments were retrieved from the Gene Expression Omnibus (GEO) and aligned using a cloud-based infrastructure. In total 137,792 samples are accessible through ARCHS4 with 72,363 mouse and 65,429 human samples. Through efficient use of cloud resources and dockerized deployment of the sequencing pipeline, the alignment cost per sample is reduced to less than one cent. ARCHS4 is updated automatically by adding newly published samples to the database as they become available. Additionally, the ARCHS4 web interface provides intuitive exploration of the processed data through querying tools, interactive visualization, and gene landing pages that provide average expression across cell lines and tissues, top co-expressed genes, and predicted biological functions and protein-protein interactions for each gene based on prior knowledge combined with co-expression. Benchmarking the quality of these predictions, co-expression correlation data created from ARCHS4 outperforms co-expression data created from other major gene expression data repositories such as GTEx and CCLE.

ARCHS4 is freely accessible at: http://amp.pharm.mssm.edu/archs4

## Introduction

The completion of the human genome project (1) enabled the quantification of mRNA expression at the genome-wide scale, initially with cDNA microarray technology (2). Genome-wide gene expression data from thousands of studies have been accumulating and made available for exploration and reuse successfully through public repositories such as the Gene Expression Omnibus (GEO) (3) and ArrayExpress (4). Since the late 1990’s software for the analysis of cDNA microarray data has matured toward established community accepted computational procedures. Now, with the advent of next generation sequencing, RNA-sequencing (RNA-seq) is becoming the leading technology for profiling genome-wide mRNA expression (Fig. 1). RNA-seq is replacing cDNA microarrays as the dominant technology due to reduced cost, increased sensitivity, ability to quantify splice variants and perform mutation analysis, improved quantification at the transcript level, identification of novel transcripts, and improved reproducibility (5).

**Fig. 1.**
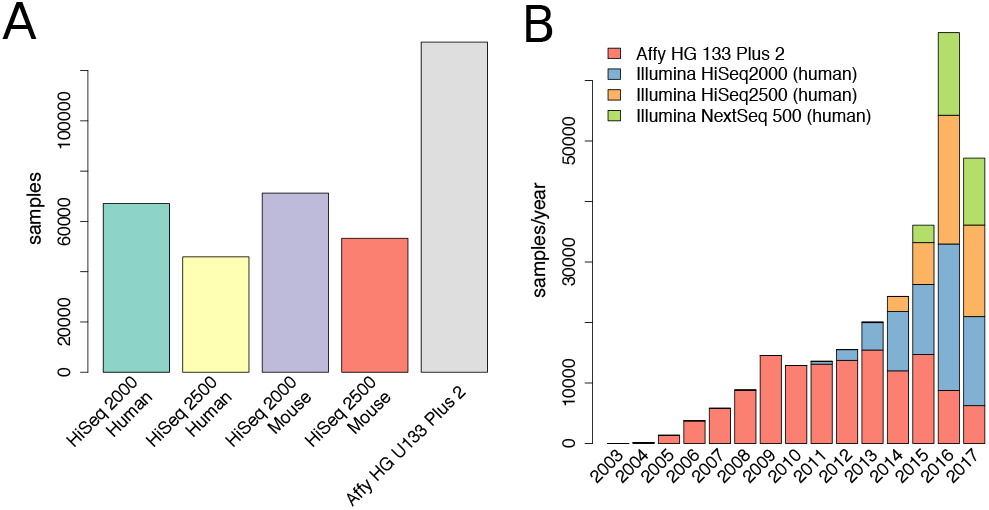
**a)** Publicly available RNA-seq samples currently available at GEO/SRA for human and mouse compared to available samples collected with the popular Affymetrix HG U133 Plus 2 platform. **b)** Total number of available RNA-seq samples collected from human cells and tissues, and those collected with Affymetrix HG U133 Plus 2 platform over time from year 2003 to 2017.

The quality of RNA-seq data depends on the sequencing depth whereby more reads per sample can reduce technical noise. Modern sequencing platforms such as Illumina HiSeq produce tens of millions of paired-end reads of up to 150 base pairs in length. The raw reads are aligned to a reference genome by mapping reads to known gene sequences. The alignment step is a computationally demanding task. The critical step of alignment of RNA-seq data is achieved by various competing alignment algorithms implemented in software packages that are continually improving (6–12). Bowtie (6) is one of the first alignment methods that gained widespread popularity. More efficient solutions were later implemented improving memory utilization with faster execution time. One of the currently leading alignment method, Spliced Transcripts Alignment to a Reference (STAR) (8), can map more than 200 million reads per hour. As a trade- off for increased computational speed, STAR requires heavy memory consumption, particularly for large genomes such as human or mouse. For mammalian genomes, STAR requires more than 30GB of random access memory (RAM). This requirement limits its application to high performance computing (HPC). This introduces a barrier for the typical experimental biologist who generates the data. Additionally, knowledge in programming, and a series of choices in regards to the alignment software, and the proper choice of input parameters to these tools, is commonly required to covert raw reads to quantified expression matrix of RNA-seq data. Retrospective analysis of large collections of previously published RNA-seq data has great promise for accelerating biological and drug discovery (13). However, many post-hoc studies rely on large datasets that are easily integrate-able into data analysis workflows whereby gene expression data is provided in processed form. For example, the Genotype-Tissue Expression project (GTEx) (14), and The Cancer Genome Atlas (TCGA) (15) RNA-seq datasets are frequently reused mainly because the data from these projects are provided in a useful processed format. GTEx currently contains 9,662 RNA-seq samples from 53 human tissues collected from >250 individuals, whereas TCGA contains at least 11,077 RNA-seq samples created from a diverse collection of patient tumors. Recent efforts such as recount2 (16) and RNAseqDB (17) attempt to simplify the access to gene expression data collected via RNA-seq to create a more unified data resource from fragmented repositories. Currently, as of mid-2017, there are more than 137,000 RNA-seq samples, collected from mammalian cells and tissues, that are accessible from the Gene Expression Omnibus (GEO) and the Sequence Read Archive (SRA); making this resource the most comprehensive repository for RNA-seq data available to date. This large collection of samples from diverse institutions, laboratories, studies and projects is much more comprehensive, but less homogeneous, compared to RNA-seq data collected for large projects such as GTEx and TCGA. However, currently, the data within GEO is provided in raw form; while the samples that have been processed, are mostly not uniformly aligned. This shortcoming makes it difficult to query and integrate this data at a global scale. To bridge the gap that currently exists between RNA-seq data generation, and RNA-seq data processing, we developed the resource all RNA-seq and ChIP- seq sample and signature search (ARCHS4). ARCHS4 provides multiple channels of processed RNA-seq data from GEO/SRA to support retrospective data analyses and reuse. ARCHS4 caters to users with different levels of computational expertise and can be employed for many post-hoc analyses and projects. The goal is to provide users with direct access to the data through a web- based user interface, while implementing a scalable and cost-effective solution for the raw data processing task. The usefulness of the resource is exemplified through case studies that show how the data assembled for ARCHS4 can be used to predict gene function and protein-protein interactions.

## Methods

### A RNA-seq Data Processing Pipeline

The RNA-seq processing pipeline employed to create the ARCHS4 resource runs in parallel on the Amazon web services (AWS) cloud. The core component of the pipeline is the alignment of raw reads to the reference genome. This process is encapsulated in deployable Docker containers (18) that currently support alignment with two leading fast aligners: STAR (8) and Kallisto (9). The memory requirement for a Kallisto Docker image is 4GB, and for STAR 30GB. All available SRA files are identified by downloading the GEO series (GSE) and GEO samples (GSM and SRA information) using the GEOquery Bioconductor package (19). Unprocessed SRA files are entered as jobs into the scheduler database of ARCHS4. The job scheduler is a dockerized web application with APIs to communicate job instructions to worker instances, and for saving the final gene count files. To allow efficient scaling of computational resources, AWS auto-scaling groups is utilized in combination with the cluster management console (ECS). For Kallisto instances, a task definition is specified running a Docker image hosted publicly at Docker Hub with 1 vCPU and a 3.9GB memory limit. The number of desired tasks specifies how many Docker images are to run in parallel. For ARCHS4, we ran 400 Docker instances of Kallisto in parallel due to AWS cap of 200 EC2 instances. The auto- scaling group is set to launch 200 m4.large general compute instances with 2 vCPUs and 8GB of memory and 200GB of SSD disk storage. Each instance is capable of running 2 Kallisto Docker instances in parallel. The alignment Docker container, once launched, continuously requests alignment jobs from the job scheduler. The job description contains the SRA file URL and the reference genome. The SRA file is downloaded from the SRA database, while Fastq-dump from SRA tools is used to detect single or paired reads file. Then, the SRA file is converted into FASTQ format. In case of a paired read file, the data is split into two FASTQ files. Kallisto or STAR alignment tools are then used to align the reads against the reference genome. The resulting output is reduced to gene counts and uploaded through the job scheduler API to the gene count database. The complete workflow is visualized as a flow chart (Fig. 2). For a subset of 1,708 FASTQ files, reads were aligned using STAR. The Docker (18) container maayan- lab/awsstar was deployed on a local Mesosphere platform (20) running on Mac Pros with 3.7 GHz Quad-Core Intel Xeon E5 and 32GB of RAM. The supported genomes are Ensemble Homo Sapiens GRCh38 with GRCh38.87 and Mus Musculus GRCm38 with GRCm38.88.

**Fig. 2.**
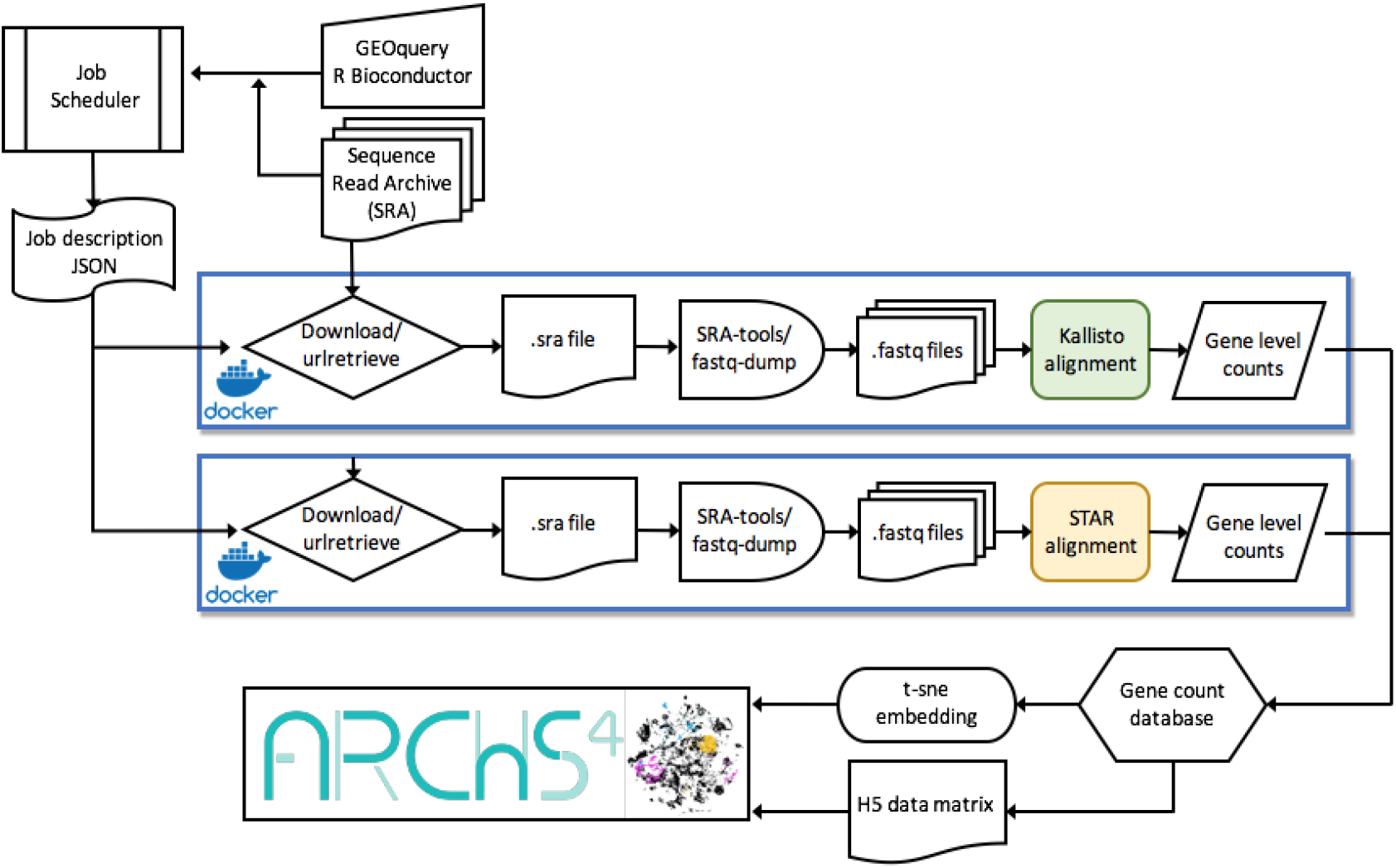
Schematic illustration of the ARCHS4 cloud-based alignment pipeline work-flow. A job scheduler instructs dockerized alignment instances that are processing FASTQ files from the SRA database in parallel. The pipeline supports the STAR and Kallisto aligners. The final results are sent to a database for post processing. Dimensionality reduction for data visualization is calculated with t-SNE and all counts are additionally stored in a H5 data matrix.

### B Post-Processing to Make the RNA-seq Data Accessible

To integrate new samples into the ARCHS4 resource, gene count files are extracted from the database and saved into a HDF5 file (21). The HDF5 files for human and mouse contain metadata describing each sample retrieved from GEO with GEOquery. The files are then deployed to Amazon S3 to be accessible for download. The 3D visualization of all samples on the ARCHS4 web-site is calculating with t-distributed stochastic neighborhood embedding (t-SNE) (22) after quantile normalization and log2 transformation of human and mouse samples separately. The t-SNE procedure uses a perplexity of 50 for the sample centric embedding, and a perplexity of 30 for the gene centric embedding using the Rtsne package in R (23). The integration of the processed data into GEO series landing pages is achieved through the ARCHS4 Chrome extension. The ARCHS4 Chrome extension, freely available at the Chrome web store, first detects whether a GEO GSE landing page is currently open in the browser. It then requests the matching GSE series matrices from ARCHS4 containing the gene expression counts and metadata information for the GSE. Additionally, the ARCHS4 Chrome extension requests JSON objects with pre-computed clustered gene expression for visualizing the samples with Clustergrammer (24). Summary statistics of sample counts and tissue specific samples are saved in a dedicated database table to be accessed by the ARCHS4 website landing page for display.

### C Sample Search with Reduced Dimensionality

To enable reliable similarity search of signatures within the ARCHS4 data matrix, the matrix is compressed into a lower dimensional representation. A projection that maintains pair-wise distances and correlations between samples is computed with the Johnson-Lindenstrauss (JL) method (25). The JL-transform reduces the original gene expression matrix *E* s *N x M* where *N* is the number of genes and M is the number samples, into a matrix 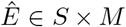, with *S < N. A* subspace of 1,000 dimensions captures the original correlation structure with a correlation coefficient of 0.99 Fig 3 a). For implementing the ARCHS4 signature search, a projection matrix *Djl* 荤 1000 x *N* is used to calculate 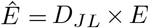. The human and mouse matrices are handled separately. For user queries, input signatures 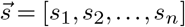 are projected onto a lower dimension 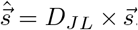. Since 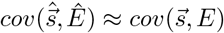, this method enables responsive real time signature similarity search with low error.

**Fig. 3.**
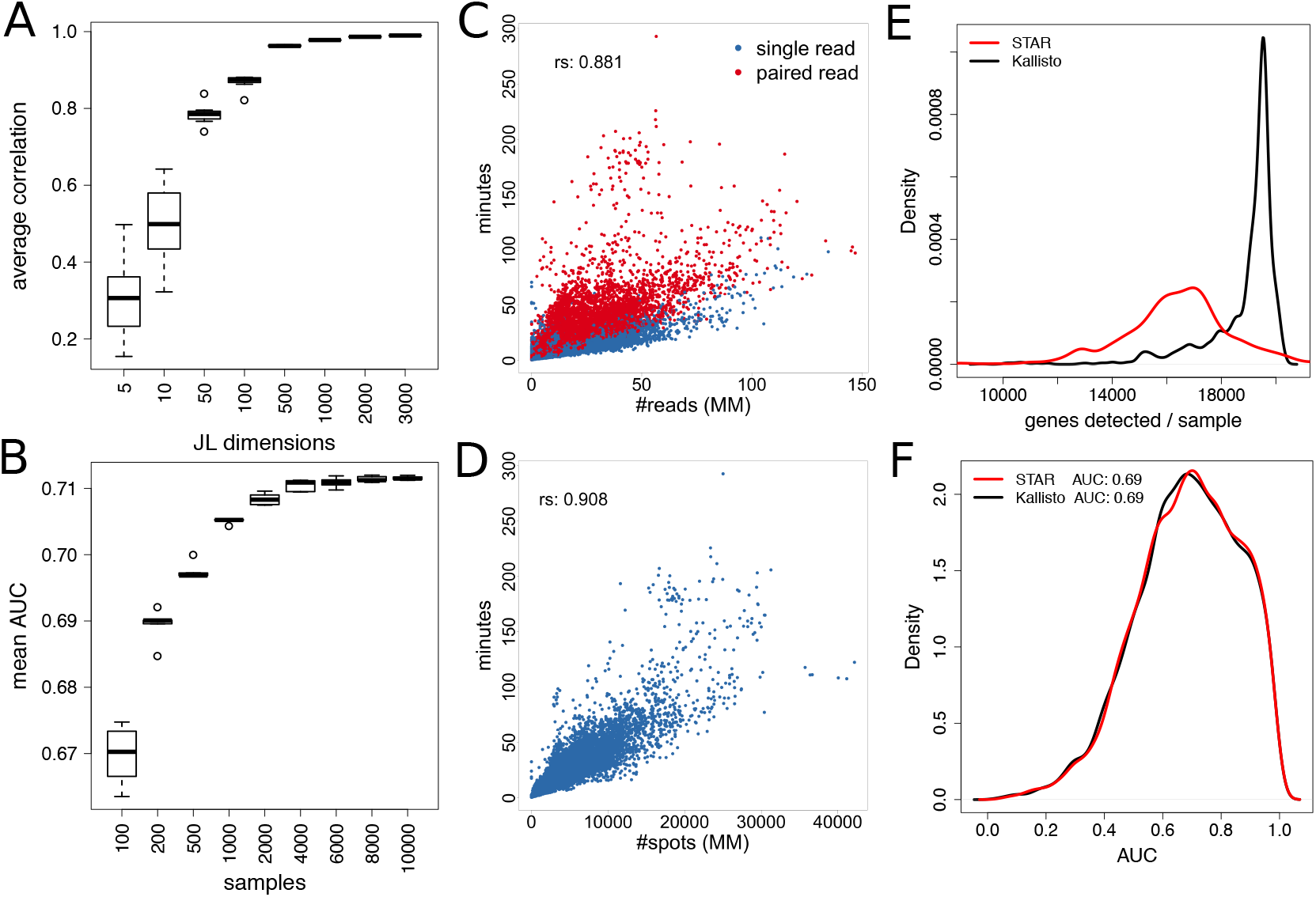
**a)** Average correlation between samples before and after applying the Johnson-Lindenstrauss dimensionality reduction. The original gene expression matrix is reduced from 34198 genes/dimensions to smaller sets of JL dimensions. For each number of JL dimensions, the procedure was repeated 10 times to obtain variances. **b)** Mean AUC for predicting GO biological processes using the ARCHS4 mouse co-expression data created from different size sets of randomly selected samples. **c)** Processing time per million reads for single read and paired end read RNA-seq for the Kallisto processing container. **d)** Elapsed time per million (MM) spots/nucleotides for completing the processing of paired read FASTQ files with the dockerized Kallisto processing container. **e)** Distribution of the number of detected genes for pipelines that utilize the Kallisto vs. STAR aligners across 1,708 randomly selected and processed human RNA-seq samples. **f)** Distribution of AUCs for predicting gene set membership for GO biological processes from co-expression matrices derived from the same set of 1,708 human RNA-seq samples processed by STAR or Kallisto aligners.

### D The ARCHS4 Interactive Web-Site

The front-end of ARCHS4 is hosted on a web server derived from the tutum/lamp Docker image which is pulled from Docker Hub. It is a web service stack running on a UNIX-based operating system with an Apache HTTP server, and a MySQL database. ARCHS4 is an AJAX application implemented with PHP and JavaScript. All visual data representations are implemented in JavaScript. The sample statistics overview of the landing page is implemented using D3.JS (26). On the data view page, the sample and gene three dimensional embedding is visualized using Three js and WebGL (27) which enable the responsive visualization of thousands of data points in 3D. Data driven queries such as signature similarity searches are performed in an R environment hosted on a dedicated dockerized Rook web server. On startup, the Rook server automatically retrieves all necessary data files, including the JT transformed gene expression table, as well as the transformation matrix, and loads them into memory for fast access from an S3 cloud repository. All Docker containers can be load balanced and run on a Mesosphere computer cluster with redundant hardware. The load balancing and port mapping is controlled through a HAProxy service. The MySQL database is hosted as a RDS amazon web service.

### E Prediction of Biological Functions and Protein-Protein Interactions

Gene-gene co-expression correlations across all human genes can be utilized to predict gene function and protein-protein interactions by exploiting the fact that genes that co-express have the tendency to also share their function and to physically interact. First, expression matrices from ARCHS4 mouse, ARCHS4 human, GTEx (14), and the cancer cell line encyclopedia (CCLE) (28) were organized into genes as the rows and samples as the columns. For the ARCHS4 data matrices 10,000 samples were randomly selected to construct gene expression correlation matrices for mouse and human separately. For GTEx and CCLE, all available samples (9,662 and 1,037 respectively) were used to build the co-expression correlation matrix for all human genes. For ARCHS4 data, functional prediction accuracy, with random subsets from the total collection, increases with the number of samples (Fig. 3b), while gains become marginal with more than 10,000 samples. Interestingly, even with a subset of 100 randomly selected samples, functional prediction accuracy is high. The quantile normalization function from the Bioconductor package preprocessCore (29) was used to normalize gene counts across samples. From the extracted expression matrices, all pairwise gene correlations were calculated. For each gene set *gsj* ∈ *GS* and each gene *gi* the mean correlation of the genes in the gene set to *gi* was calculated. Self- correlations when *g_i_* ∈ *gsj* were excluded. Hence, the resulting gene set membership prediction matrix *GM* ∈ *M* × *N* for *M* genes and *N* gene sets is generated by the following procedure:

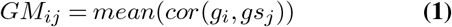

*GMi* is then sorted from high to low based on the correlation levels. For each row *i* within *GM*, a vector 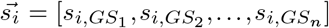 is then constructed where *si,GSj* ∈ {0,1} and *s_i_,gsj* is 1 if gene *gi* is already known to be in the gene set *GS*. This vector is sorted and used to compute the area under the curve (AUC) from the cumulative sum of 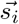 using trapezoidal integration. To predict protein-protein interactions (PPI), the three PPI networks: hu.MAP (30), BioGRID (31) and BioPLEX (32) are first converted to a gene set libraries as described in (33). Then, to predict PPIs, the same procedure for functional predictions was applied.

## Results

### F The ARCHS4 Web-Site

The ARCHS4 website supports multiple complementary ways of accessing the processed RNA- seq gene expression data. For programmatic access, the download section provides users with the ability to download all the gene expression data for human and mouse in H5 format. The H5 files contain extensive metadata information retrieved from GEO. This metadata can be queried to extract samples of interest by a key term. Additionally, programmatic access to ARCHS4 supports exploration of gene expression matrices of interest through multiple search functions. The ARCHS4 website visualizes all the processed samples, and alternatively all human or mouse genes, as interactive 3D t-SNE plots. In the sample centric view, all samples can be searched based on metadata terms. ARCHS4 performs a string text search of the GEO metadata to retrieve samples by matching the string terms. For example, searching Pancreatic Islet in the human context will return 1,829 samples from 10 independent GEO series. After the search is complete, the samples are highlighted in the 3D visualization, and an auto-generated R script is provided for download. Executing the R script builds a local expression matrix in TSV format with the samples as columns and the genes as the rows. The signature search in ARCHS4 enables searching samples by data, matching high and low expressed genes from input sets to high and low expressed genes across all ARCHS4 processed samples. Signature similarity is approximated via the Johnson-Lindenstrauss (JL) (25) transformed gene expression space. Under the enrichment search tab, samples can be selected by precomputed enrichment for annotated gene-sets. Gene set libraries from which annotated gene sets are currently derived from are: ChEA (34), ENCODE (35), (36), Gene Ontology (GO) (37), KEGG pathways (38), and MGI mammalian phenotype (39). The ARCHS4 three dimensional viewer also supports manual selection of samples through a snipping tool, whereas the colors used to highlight samples can be changed by the user. The gene-centric view of ARCHS4 provides the same manual selection feature as with the sample view. Selected gene lists can be downloaded directly from the ARCHS4 website or sent to Enrichr (40, 41), a gene set enrichment analysis tool, for further functional exploration. Gene sets can be highlighted in the 3D view. Additionally, individual genes can be queried to locate ARCHS4 gene landing pages. These single gene landing pages contain predicted biological functions based on correlations with genes assigned to GO categories; predicted upstream transcription factors based on correlation with identified putative targets from ChEA and ENCODE; predicted knockout mouse phenotypes based on annotated MGI mammalian phenotypes; predicted human phenotypes based on co- expression correlation with genes that have assigned human phenotypes in the human phenotype ontology (42); predicted upstream protein kinases based known kinase-substrates from KEA, and membership in pathways based on co-expression with pathway members from KEGG. The single gene landing pages also list the top 100 most co-expressed correlated genes for each individual gene. Additionally, for 53 distinct tissues, tissue expression levels are visualized for each gene. The tissue expression display is visualized as a hierarchical tree with tissues grouped by system and organ. Similarly, cell line gene expression profile for 67 widely used cell lines across tissue types can be accessed through the gene landing pages of ARCHS4. ARCHS4 processed data can be accessed via the ARCHS4 Chrome extension, which is freely available from the Chrome web store. The Chrome extension detects GEO series landing pages and inserts a Series Matrix File (SMF) for download for each series that have been processed by the ARCHS4 pipeline. Each SMF contains read counts for all available samples in the series. The sample expression is also visualized as a heatmap using the Clustergrammer plugin (24). Clustergrammer loads JSON files containing the z-score normalized gene expression of the top 500 most variable genes across the series, and embeds the interactive heatmap directly into the GEO series landing page.

### G RNA-seq Alignment Pipeline Speed and Cost

The pipeline speed is measured by the elapsed time from job submission until completion for 31,825 samples processed from the GEO/SRA database. This includes: the SRA file download, FASTQ file extraction from the SRA data format, alignment to the reference genome, mapping the transcript counts to the gene level, and writing the final result to the database. Processing time of a single FASTQ file is on average 11 minutes. For samples by number of spots and single and paired end samples, the benchmark is applied using Amazon EC2 on-demand m4.large instances with 8GB of memory and 2 vCPUs. Instances running with 200GB of hard-drive storage. Each instance can run 2 dockerized alignment pipeline containers in parallel. At the time of the benchmark, the cost of the on demand m4.large instances was $0.1/h. This results in an average compute cost for one processed SRA file to be $0.00917. The alignment time correlates with the number of reads (Spearman correlation coefficient r=0.881) and the processing time increases linearly with the number of spots aligned with some variance (*r* = 0.901, paired reads) due to performance differences between cloud computing instances (Fig. 3cd). Paired end read RNA-seq experiments require more time during the alignment process due to the increased number of spots that have to be processed. The ARCHS4 pipeline is to our knowledge, the most cost effective cloud based RNA-seq alignment infrastructure published to date.

### H STAR vs. Kallisto Comparison

To achieve its fast and cost effective solution, ARCHS4 utilizes the Kallisto aligner (9) to process all samples. However, it is not clear whether the improved speed and cost provided by Kallisto comes with the cost of drop in output quality. To benchmark Kallisto, a random subset of 1,708 human samples processed by ARCHS4 were also aligned with STAR (8). While Kallisto and STAR return similar gene expression profiles, there are profound differences between the output produced by the two algorithms. In general, Kallisto detects more genes than STAR (Fig. 3ef). The average Pearson correlation of the z-score transformed samples between the Kallisto and STAR output is 0.77. However, the number of detected genes does not directly translate to qualitative advantage of Kallisto over STAR. To test the quality of the generated gene expression matrices and their gene correlation structure, the ability to predict GO biological processes for single genes, as described in detail in the methods section, was utilized as a benchmark. The co-expression matrices used for predicting GO biological processes were created from the data aligned by Kallisto or STAR. The quality of the predictions is almost identical with an average area under the curve (AUC) of 0.69 for predictions made by processed data from both sources (Fig. 3f).

### I Read Quality across Institutions

The percentage of aligned reads over total reads for each FASTQ file varies significantly across labs, projects, and sequencing cores due to various reasons. Since each sample from GEO/SRA is annotated with the producers of the data, the percent of aligned reads by institution can be plotted (Fig. 4). The 18 institutions that so far produced more than 100 unique samples from more than 10 gene expression series of human RNA-seq samples available on the GEO/SRA database show that the highest percentage of successful aligned reads is by the University of Utah with a median of 84%. The 341 samples that originated from the University of Utah come from 14 distinct gene expression series. It should be noted that observed differences in the fraction of aligned reads is not necessarily an indicator of the performance of the sequencing core within an institution, but can be attributed to the quality of the samples. For example, samples from formalin fixated tissues will suffer from RNA degradation which will result in lower percent of aligned reads. On average, 63% of reads were aligned across all 65,429 the processed human RNA-seq samples, whereas 59% of all mouse RNA-seq reads from 72,363 samples were aligned to matching genes.

**Fig. 4.**
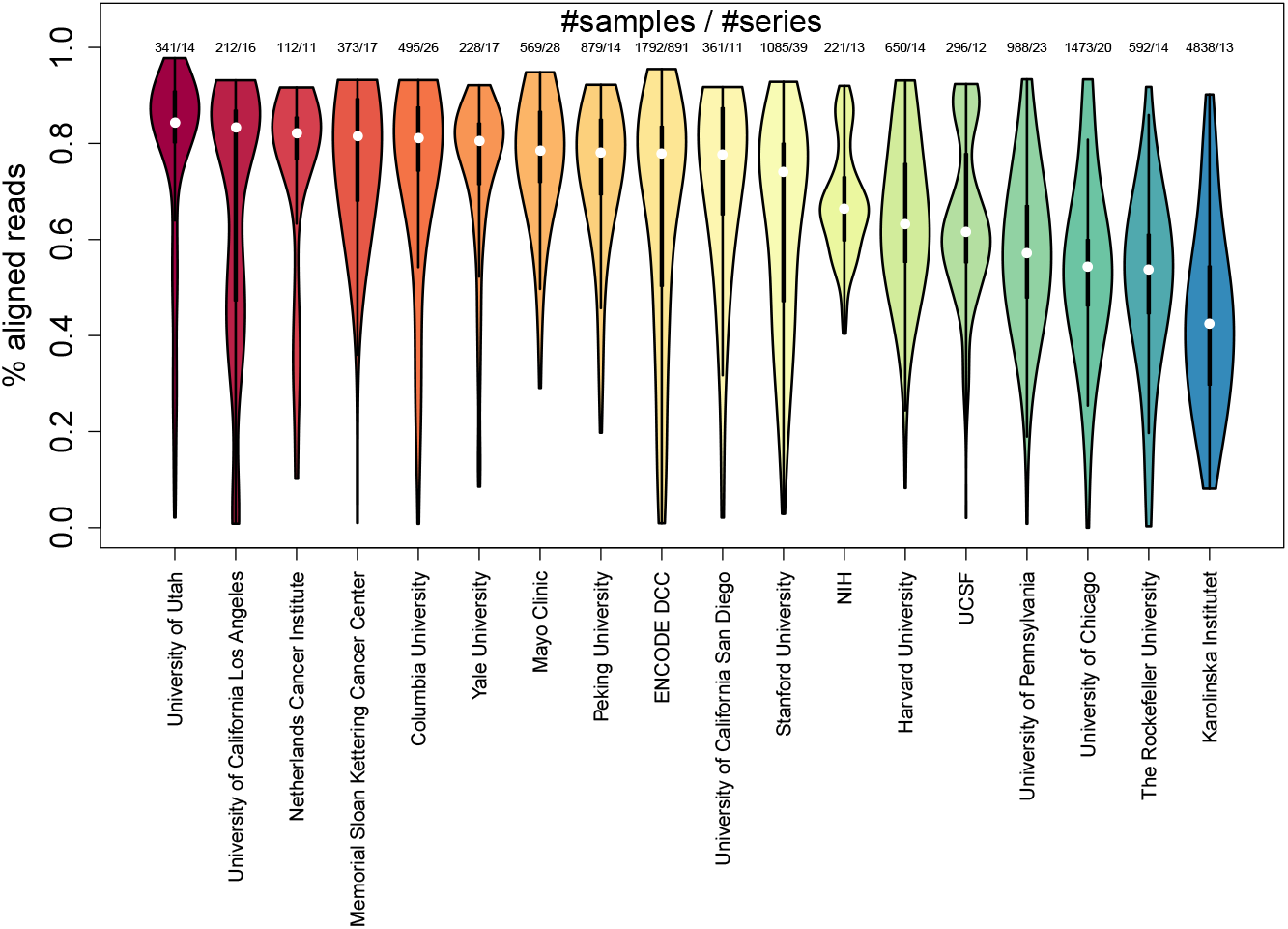
Distribution of the percentage of aligned reads from human RNA-seq samples that are successfully aligned with Kallisto by institution as it is reported at the GEO submission pages. The selected institutions that are shown have more than 100 samples from more than 10 different gene expression series.

### J Prediction of Gene Function and Protein-Protein Interactions with ARCHS4 Data

Gene function and protein-protein interactions can be potentially predicted using co-expression data, whereas the data that is processed for ARCHS4 provides a rich resource for generating gene co-expression networks. Evaluating the quality of co-expression networks to predict protein interactions and biological functions can also provide an unbiased benchmark to compare the ARCHS4 resource to other major RNA-seq and microarray repositories. The hypothesis that gene function and protein interactions can be predicted using co-expression data implies that co- expressed genes tend to share their function and physically interact. In this process, genes are assigned predicted biological functions only when they are highly correlated to genes within a set of genes already annotated to have the same biological function. Similarly, a gene product is predicted to interact with another protein if the known direct protein interactors for that other protein are highly co-express with the gene product protein. We evaluate the human and mouse ARCHS4 datasets by comparing them to co-expression matrices created in the same way from the CCLE and GTEx resources. All gene expression datasets produce, on average, significant ability to predict both biological functions and protein interactions. This suggests that gene expression correlations derived from large scale expression datasets are predictive of biological function and protein interactions. In almost all the tested categories, the ARCHS4 mouse and human datasets outperformed the predictions made with co-expression data created from the CCLE and GTEx datasets significantly (Table 1). The most accurate predictions for GO biological processes, GO molecular functions, KEGG pathways, Human Phenotype Ontology terms, predicted upstream kinases, and MGI Mammalian Phenotype terms are achieved with the ARCHS4 mouse gene co- expression data followed by the ARCHS4 human data. The co-expression data from GTEx outperforms the co-expression data created from the CCLE for GO biological processes and the phenotype libraries, whereas the predictability GTEx data is lower than CCLE for the upstream regulatory transcription factor predictions. P-values are calculated for the mean between methods. For the ARCHS4 mouse co-expression data the area under the curve (AUC) distributions for predicting gene function are significant across all categories, but most successful in predicting GO biological processes with median AUC of 0.745, and membership in KEGG pathways with a median AUC of 0.797 (Fig. 5a). While predicting protein function with co-expression data has been attempted successfully by many before, it is less established whether co-expression data can be used to also predict protein- protein interactions (PPI). A similar strategy was employed to predict PPI using prior knowledge about PPI from three unique PPI resources: hu.MAP (30), BioGRID (31) and BioPLEX (32). These three PPI resources are unique in the following way: the PPI from BioGRID are the composition of interactions from thousands of publications that report only few interactions; the PPI from BioPLEX are bait-prey interactions from a massive mass-spectrometry experiment; whereas hu.MAP is made of data from three mass-spectrometry experiments integrated with sophisticated computational methods that also consider prey-prey interactions to boost interaction confidence. Importantly, none of these three resources utilize knowledge from mRNA co- expression data to confirm PPI. The overlap of shared interactions between the three PPI networks is relatively low with hu.MAP and BioPLEX sharing more than 10% of their interaction (Fig. 5b). This is likely because a part of BioPLEX is contained within hu.MAP. Predicting PPI using knowledge from these three PPI resources, the ARCHS4 mouse co-expression data is the most predictive with a median AUC of 0.85, 0.66 and 0.64 for hu.MAP, BioGRID and BioPLEX respectively (Fig. 5c).

**Fig. 5.**
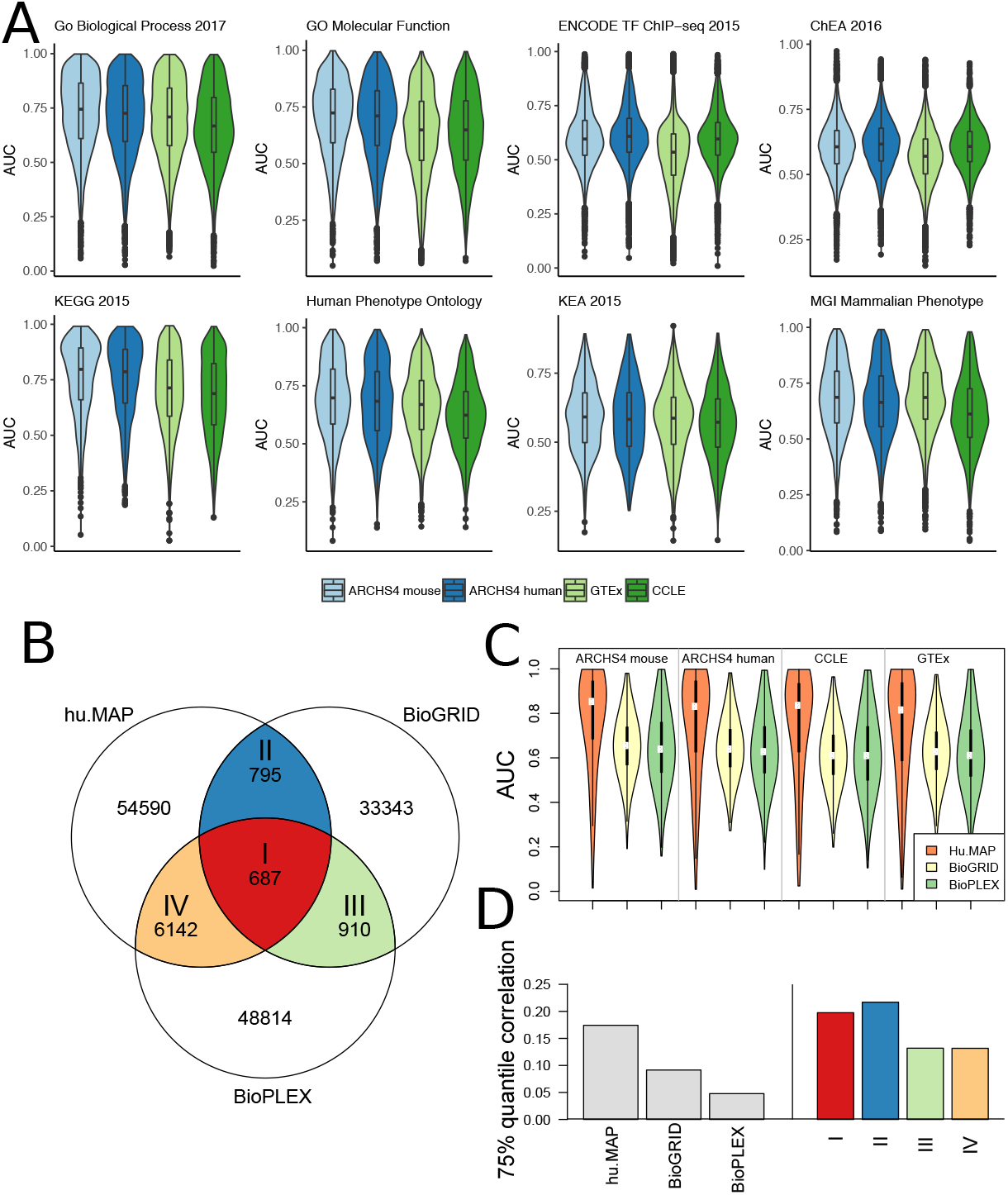
**a)** The distribution of AUC for gene set membership prediction of gene annotations from eight gene set libraries with co-expression data created from ARCHS4 mouse, ARCHS4 human, GTEx and CCLE. The gene set libraries used to train and evaluate the predictions are ChEA, ENCODE, GO Biological Process, GO Molecular Function, KEA, KEGG Pathways, Human Phenotype Ontology, and MGI Mammalian Phenotype Level 4. These libraries were obtained from the Enrichr collection of libraries. **b)** Venn diagram showing the intersection of edges between three PPI databases hu.MAP, BioGRID and BioPLEX. **c)** Distribution of AUC for protein-protein interaction prediction from gene co-expression data created in the same way from ARCHS4 mouse, ARCHS4 human, CCLE and GTEx. **d)** Bar plot of the pairwise correlation between genes with reported protein-protein interactions for the three PPI networks hu.MAP, BioGRID and BioPLEX in ARCHS4 mouse expression. The right tail of the gene pair correlation distribution of is shown by the 75% quantile.

**Table 1.** Comparison of functional prediction for ARCHS4 mouse and human gene expression compared to GTEx and CCLE. Δ median is the difference in median AUC between ARCHS4 mouse and the other data sets. The significance of difference of the mean is calculated by t-test for observed AUC distributions. The highest median per gene set library are highlighted in bold.

| Gene Set Library |  | ARCHS4 mouse | ARCHS4 human | GTEx | CCLC |
| --- | --- | --- | --- | --- | --- |
| Go Biological Process 2017 | median | **0.745** | 0.726 | 0.709 | 0.667 |
| | $\Delta$ median | 0 | -0.0186 | -0.0356 | -0.0773 |
|  | p-value | 1 | 7.70E-08 | 1.47E-27 | 8.50E-124 |
| GO Molecular Function | median | **0.724** | 0.710 | 0.649 | 0.649 |
| | $\Delta$ median | 0 | -0.0134 | -0.0752 | -0.0752 |
|  | p-value | 1 | 0.0174 | 1.08E-78 | 7.93E-64 |
| ENCODE TF ChIP-seq 2015 | median | 0.596 | **0.608** | 0.5349 | 0.596 |
| | $\Delta$ median | 0 | 0.0124 | -0.0610 | 0.000271 |
|  | p-value | 1 | 5.54E-13 | 0 | 0.000476 |
| ChEA 2016 | median | 0.606 | **0.617** | 0.570 | 0.608 |
| | $\Delta$ median | 0 | 0.0104 | -0.0363 | 0.00139 |
|  | p-value | 1 | 1.95E-17 | 5.44E-266 | 0.758 |
| KEGG 2015 | median | **0.797** | 0.786 | 0.713 | 0.688 |
| | $\Delta$ median | 0 | -0.0109 | -0.0838 | -0.109 |
|  | p-value | 1 | 0.210 | 5.76E-20 | 2.56E-35 |
| Human Phenotype Ontology | median | **0.698** | 0.683 | 0.669 | 0.623 |
| | $\Delta$ median | 0 | -0.0144 | -0.0284 | -0.0745 |
|  | p-value | 1 | 0.00251 | 2.38E-10 | 6.05E-48 |
| KEA 2015 | median | **0.591** | 0.583 | 0.587 | 0.572 |
| | $\Delta$ median | 0 | -0.00880 | -0.00439 | -0.0190 |
|  | p-value | 1 | 0.431 | 0.0459 | 0.00365 |
| MGI Mammalian Phenotype | median | **0.687** | 0.6639 | 0.686 | 0.612 |
| | $\Delta$ median | 0 | -0.0227 | -0.000726 | -0.0749 |
|  | p-value | 1 | 3.83E-08 | 0.537 | 9.97E-83 |

The fact that PPI from hu.MAP can be predicted at a much higher accuracy compared to the other two networks suggests that protein-protein interactions from hu.MAP tend to have higher pairwise mRNA co-expression correlations. The 75%-quantile of interaction correlation in hu.MAP is 0.174 compared to BioGRID (0.0915) and BioPLEX (0.0478); whereas the intersections between the PPI networks (I, II, III, IV) tend to have a higher 75%-quantile of correlations with 0.198, 0.217, 0.131 and 0.132 suggesting that aggregating evidence from experiments that detect PPI, is most likely boosting confidence of real interactions (Fig. 5d). This also further supports that mRNA co-expression data can be used to predict PPI. The predicted PPI and predicted biological functions provide a plethora of computational hypotheses that could be further validated experimentally.

## Discussion and Conclusion

The ARCHS4 resource of processed RNA-seq data is created by systematically processing publicly available raw FASTQ samples from GEO/SRA. This resource can facilitate rapid progress of retrospective post-hoc focal and global analyses. The ARCHS4 data processing pipeline employs a modular dockerized software infrastructure that can align RNA-seq samples at an average cost of less than one cent (US $0.01). To our knowledge, this is an improvement of more than an order of magnitude over previously published solutions. The automation of the pipeline enables constant updating of the data repository by regular inclusion of newly published gene expression samples. The pipeline is open source and available on GitHub so it can be continually enhanced and adopted by the community for other projects. The pipeline uses Kallisto as the main alignment algorithm which was demonstrated through an unbiased benchmark to perform as well as, or even better than, another leading aligner, STAR. We compared the ARCHS4 co-expression data with co-expression data we created from other existing gene expression resources, namely GTEx and CCLE, and demonstrated how co-expression data from ARCHS4 is more effective in predicting biological functions and protein interactions. This could be because the data from ARCHS4 is more diverse. The fact that the data within ARCHS4 is from many sources has its disadvantages. These include batch effects and quality control inconsistencies. Standard batch effect removal methods are not applicable to the entire ARCHS4 data, but may useful for improving the analysis of segments of ARCHS4 data. The ARCHS4 web application and Chrome extension enable users to access and query the ARCHS4 data through metadata searches and other means. For data driven queries, the unique JL dimensionality reduction method is implemented to maintain pairwise distances and correlations between samples even after reducing the number of dimensions by two orders of magnitude. Reducing the data to a lower dimension facilitates data driven searches that return results instantly. The gene expression data provided by ARCHS4 is freely accessible for download in the compact HDF5 file format allowing programmatic access. The HDF5 files contain all available metadata information about all samples, but such metadata can be improved but having it follow naming standards and linking it to biological ontologies. The ARCHS4 three dimensional data viewer lets users gain intuition about the global space of gene expression data from human and mouse at the sample and gene levels. The interface supports interactive data exploration through manual sample selection and highlighting of samples from tissues and cell lines. With the available data, we constructed comprehensive gene landing pages containing information about predicted gene function and PPI, co-expression with other genes, and average expression across cell lines and tissues. For a variety of tissues and cell lines, gene expression distributions are calculated for each gene. Such data can complement tissue and cell line expression resources such as BioGPS (43) and GTEx (14) as well as resources that provide accumulated knowledge about genes and proteins such as GeneCards, the Harmonizome and the NCBI gene database (44–46). Overall, the ARCHS4 resource contain comprehensive processed mRNA expression data that can further enable biological discovery toward better understanding of the inner-workings of mammalian cells.

## ACKNOWLEDGEMENTS

This work is partially supported by the National Institutes of Health (NIH) grants U54HL127624 and U54CA189201, as well as cloud credits from the NIH BD2K Commons Cloud Credit Pilot project to AM.

